# Fine-scale structure of a whole regional population through genetics and genealogies

**DOI:** 10.1101/2025.09.23.678060

**Authors:** Gilles-Philippe Morin, Claudia Moreau, Amadou Barry, Simon L. Girard

## Abstract

The Saguenay–Lac-Saint-Jean (SLSJ) region of Quebec, Canada, a population shaped by a prominent founder effect, has long been considered genetically homogeneous. This study comprehensively investigates the fine-scale population structure within SLSJ by integrating genotype data from the CARTaGENE cohort with extensive genealogical records from the BALSAC population register. A time-efficient algorithm was developed to compute billions of kinship coefficients from genealogies in order to analyse an entire generation. We demonstrate a striking concordance between realised (genetic) and expected (genealogical) kinship (r = 0.78). From both kinship measures, we reveal fine-scale population structure at the municipal level within SLSJ, challenging the notion of regional homogeneity. Our analysis highlights an east–west genetic gradient and uncovers migratory streams and differential founders’ genetic contributions that shaped the genetic landscape of this population. This research provides insights into the interplay of genetics, demography, and historical events, underscoring the importance of fine-scale population structure in genetic studies and reaffirming the power of large-scale genealogical data.

## INTRODUCTION

The success of genetic association studies, such as genome-wide association studies (GWAS), depends on understanding and controlling for bias and confounding factors, including population structure^1,2^. Such structure influences the distribution of genetic variants^3,4^, including those associated with disease^5–7^. Invaluable insights were obtained by studying the fine-scale population structure of founder populations, revealing how historical migration events and geographical barriers have shaped the genetic landscape of contemporary populations^8^. In such populations, a smaller effective population size is expected to lead to homogeneity through increased homozygosity and reduced genetic diversity.

The Saguenay–Lac-Saint-Jean (SLSJ) region of Quebec, Canada, represents a remarkable model for population genetics research due to its well-documented founder effect, recent settlement and unique demographic history. From the seventeenth century, European settlers of mostly French origin have migrated along the Saint Lawrence River, then founded the Charlevoix region, before settling in Saguenay and Lac-Saint-Jean from 1838^9^. This serial founder effect was accentuated by a 25-fold population surge in SLSJ within a century, mainly due to high birth rates^10,11^. The region’s genetic characteristics are documented through the BALSAC population register^12^, a comprehensive database documenting civil records, primarily Catholic marriages, of French-Canadian individuals from the 1660s (New France) to the 1960s (Quebec), encompassing over three centuries of demographic history. This demographic history led to several variants being more frequent in the region^5^ and to the increase of some rare diseases^13–16^. Genetic testing and screening is currently available^14,17^ for six of these diseases, highlighting the clinical relevance of understanding this population’s genetic structure. Some researchers have characterized the genetic background of SLSJ as relatively homogeneous^14,18^, suggesting that, unlike the broader provincial context, the region might lack meaningful structure. Yet, demographic studies indicate that population structure does exist within SLSJ^19,20^, challenging the assumption of homogeneity.

Previous research in the French-Canadian population has shown that both genotype and genealogical data correlate substantially^8,21–23^. This shows that genealogy may be used as a proxy to study the genetic structure of populations without being restricted to genotyped individuals, who are often originating from urban areas. Yet, despite the wealth of genealogical and demographic data available for the SLSJ region, significant challenges remain in adequately visualising the region’s population structure at a large scale. Traditional methods for analysing population structure, while effective for smaller samples, face computational limitations when applied to entire populations or large generational cohorts. Previous studies have used kinship coefficients derived from genealogical data to visualise population structure^21,24^, but the computational expense of applying these calculations to whole populations using available software has been prohibitive. This limitation primarily stems from algorithms that repeatedly compute kinship for redundant pairs of individuals within a genealogy^25^, making analysis of complete populations computationally intractable. Although recent efforts have optimized these computations^26,27^, implementing these methods at the scale needed for whole-population analysis—particularly for visualization purposes—has remained challenging, restricting samples to hundreds or a few thousand individuals at a time. The absence of appropriate analytical methods has thus limited comprehensive investigation of fine-scale structure in populations like SLSJ.

This study aims to comprehensively investigate the fine-scale population structure within the SLSJ region by integrating genotype and genealogical datasets. The primary objective of this research is to compare realised kinship, calculated from genetic identity-by-descent (IBD) segments, and expected kinship, inferred from detailed genealogical records using a hybrid algorithm. We also retrace the population’s origins through the expected genetic contribution of the region’s founders. By doing so, this article reveals a fine-scale population structure shaped by geographical boundaries, differential founders’ genetic contribution and socioeconomic factors.

## SUBJECTS AND METHODS

This study was approved by the ethics board of the Université du Québec à Chicoutimi (UQAC).

### Genotype data and cleaning

Genotype data was drawn from the CARTaGENE cohort^28^, which comprises 43,032 participants aged 40-69 (in 2005) recruited in the main urban areas of the Quebec Province, notably in the city of Saguenay.

Of those participants, 29,337 individuals were genotyped using a variety of platforms, including Omni 2.5, GSAv1 with a multi-disease panel, GSAv1, GSAv2 with a multi-disease panel, GSAv3 with a multi-disease panel, GSAv2 with a multi-disease panel and add-on (see https://cartagene.qc.ca/files/documents/other/Info_GeneticData3juillet2023.pdf for details). Each dataset was processed independently using PLINK v1.9^29^, retaining individuals with at least 95% genotyping across all SNPs. At the SNP level, we retained autosomal variants with a call rate of at least 95% across individuals and that conformed to Hardy–Weinberg equilibrium (*p* > 10^−6^, calculated within each dataset). All chips were merged, and the final dataset comprised 148,200 SNPs across 28,358 individuals.

### Genealogical data

Genealogical data was obtained from the BALSAC population register^12^. Two different genealogical datasets have been extracted for this study: First, 9,405 Quebec residents from the CARTaGENE cohort have been matched to BALSAC records. Second, to analyse a whole generation, we also selected 80,348 probands (i.e. non-parent individuals) married in SLSJ from 1931 to 1960. For both probands’ sets, a multigenerational genealogy extending up to 19 generations was reconstructed. When available, information on the municipality, region, and year of marriage was included; to ensure confidentiality, marriage years are rounded up to the nearest 5. Individual anonymity is rigorously maintained through the use of unique, non-identifying codes.

Traditional geographic subdivisions split SLSJ into Lower Saguenay, Upper Saguenay, and Lac-Saint-Jean^19^. However, it has been demonstrated that geographic features and natural barriers, such as watersheds, can influence patterns of population structure^8^. We carefully carved eight subdivisions along primary watercourses to consolidate smaller communities and municipalities were grouped based on these watercourse boundaries (Supplementary Table 1 and Supplementary Fig. 1). On average, we observed reduced migration in the last generation within our defined geographical subdivisions than between them (Supplementary Fig. 2).

### IBD sharing (realised kinship)

Pairwise IBD segments were inferred on phased genotypes using Refined IBD (version 17Jan20)^30^ and Beagle (version 18May20)^31^. A matrix of realised kinship estimation was computed by summing shared segment lengths of at least 2 centiMorgans (cM) using a Python 3 ported version of `relatedness_v1.py` from Sharon R. Browning (see https://faculty.washington.edu/sguy/ibd_relatedness.html). The diagonal was set to 1.

### Genealogical kinship coefficients (expected kinship)

Expected kinship (*ϕ*)^32^ was used to infer and visualise population structure through genealogical data. It corresponds to the probability that, at one locus, one randomly picked allele from individual *i* and one randomly picked allele from individual *j* are IBD. Expected kinship coefficients were computed between all pairs of probands from whole SLSJ and CARTaGENE genealogical datasets. Of the 80,348 SLSJ probands, 77,088 have a genealogy deemed sufficiently complete, i.e. they are related (*ϕ* > 0) with at least half of the probands, and the others were removed. Of the remaining individuals, 26,445 are non-siblings (*ϕ* > 0.2). Of the 9,397 probands in CARTaGENE data, 7,970 are non-siblings, and have a genealogy deemed sufficiently complete using the same criterion.

#### Hybrid algorithm for time-efficient computation of billions of kinship coefficients

To efficiently compute kinship coefficients across all pairs of individuals, we implemented a hybrid of two existing algorithms. Karigl’s algorithm^25^ estimates kinship coefficients on a per-proband-pair basis, enabling parallel processing of the kinship matrix, as in the GENLIB package for R^33^. However, it reprocesses each ancestor every time they appear, either multiple times across genealogies (kinship) or repeatedly within the same genealogy (inbreeding), which makes it computationally costly. Kirkpatrick’s algorithm^27^ addresses this inefficiency by partitioning the genealogy into successive sub-genealogies (e.g., one generation at a time) and computing kinship coefficients in a top-down manner, with the probands of one sub-genealogy treated as the founders of the next, thereby minimizing redundant ancestor processing. Yet, Kirkpatrick’s approach is not designed for parallel execution. Our method combines the strengths of both: we divide the genealogy into generational sub-genealogies, as in Kirkpatrick’s approach, but compute each sub- genealogy’s kinship matrix in parallel, achieving substantial time gains. For instance, computing kinship for all 6,455,801,104 pairs of 1931–1960 BALSAC SLSJ probands required only two minutes with our algorithm compared to more than a week for the existing R GENLIB package.

Our new implemented algorithm is provided as part of GeneaKit, which is freely available at https://github.com/Genopop/geneakit under the MIT licence, using the `phi()` function for kinship coefficients. A pseudocode of the algorithm is available in the supplementary material (Supplementary Text 1).

### Expected genetic contributions

We also computed the expected genetic contribution of the region’s founders as a way to measure the region’s migratory origins. Region’s founders are defined as individuals who married in the region whereas their parents married elsewhere. This computation was done using GeneaKit’s `gc()` function, from all region’s founders to all 80,348 genealogical probands who married between 1931 and 1960 in the SLSJ region. The expected genetic contribution of an ancestor corresponds to the probability that an allele is passed down to a given descendant. A founder’s region of origin is defined as the region of marriage of their parents.

### Clustering and visualisation

CARTaGENE individuals originate from all regions of Quebec, it is thus necessary to identify those who are most likely to originate from SLSJ. Uniform Manifold Approximation and Projection (UMAP)^34^ was performed on both the realised and expected kinship matrices, using precomputed distances of (*1 – ϕ*). UMAP was run with 10 components, 15 neighbors, a minimum distance of 0.0, and a fixed random seed (random state = 0), to maximise clustering and reproducibility. Hierarchical Density-Based Spatial Clustering of Applications with Noise (HDBSCAN) from `scikit-learn`35 was then fitted to the UMAP outputs to identify individuals most likely originating from the SLSJ region, using a `min_cluster_size` of 25 and a `cluster_selection_epsilon` of 0.3. Those parameters were chosen as they allowed separation of regional populations in a previous study using the same cohort^28^. From the clustering results for both the realised and expected kinship matrices, the cluster containing the highest number of individuals of known SLSJ origin (i.e. whose four grandparents married in the region) was identified. 597/1091 individuals in the realised kinship cluster and 574/1022 individuals in the expected kinship cluster are of known SLSJ origin. The final set of 904 individuals was then determined as the intersection of these two ‘best’ clusters (one from the realised kinship matrix clustering and one from the expected kinship matrix clustering). Of those 904 individuals, 564 are of known SLSJ origin. For visualisation, UMAP was applied to the kinship matrices using their precomputed distances (*1 – ϕ*), 2 components, and a minimum distance of 0.5.

## RESULTS

### Comparison between genetic and genealogical structure

The CARTaGENE cohort comprises individuals with both genotype and genealogical data, of which 8,069 probands have sufficient genealogical completeness (see methods). UMAP projections based on realised and expected kinship display strikingly similar population structure patterns (Fig. 1); a linear regression between realised and expected kinship further supports this consistency with a Pearson correlation coefficient of 0.78 (*p* < 0.001). HDBSCAN clustering analysis identified a subset of 904 individuals most likely originating from the SLSJ region (Supplementary Fig. 3). Within this subset, UMAP projections derived from both realised and expected kinship (Fig. 2) reveal partially overlapping gradients of individuals from the diverse subdivisions of SLSJ (Supplementary Fig. 4).

**Fig. 1:**
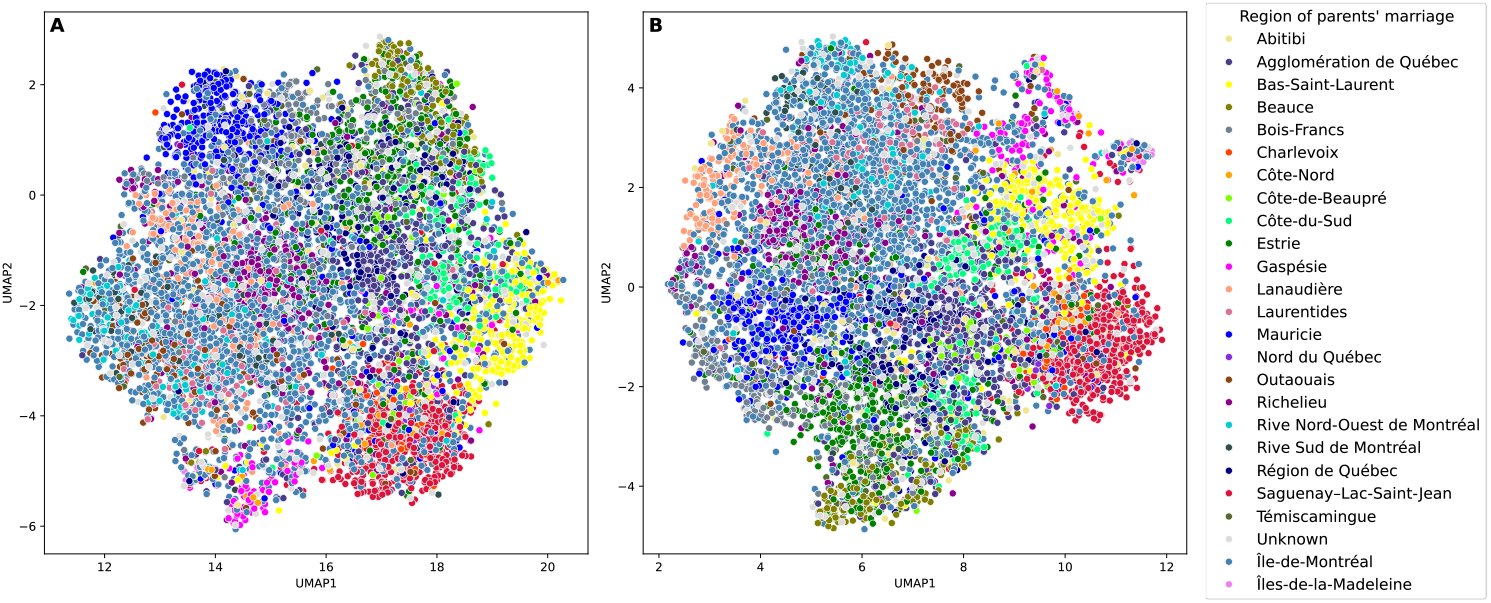
UMAP of CARTaGENE individuals based on pairwise realised (A) and expected (B) kinship. UMAP projection of 7,970 individuals from the CARTaGENE cohort, computed from their (*A*) realised kinship and (*B*) expected kinship transformed as a precomputed distance (*1 – ϕ*). The colours represent the region of parents’ marriage.

**Fig. 2:**
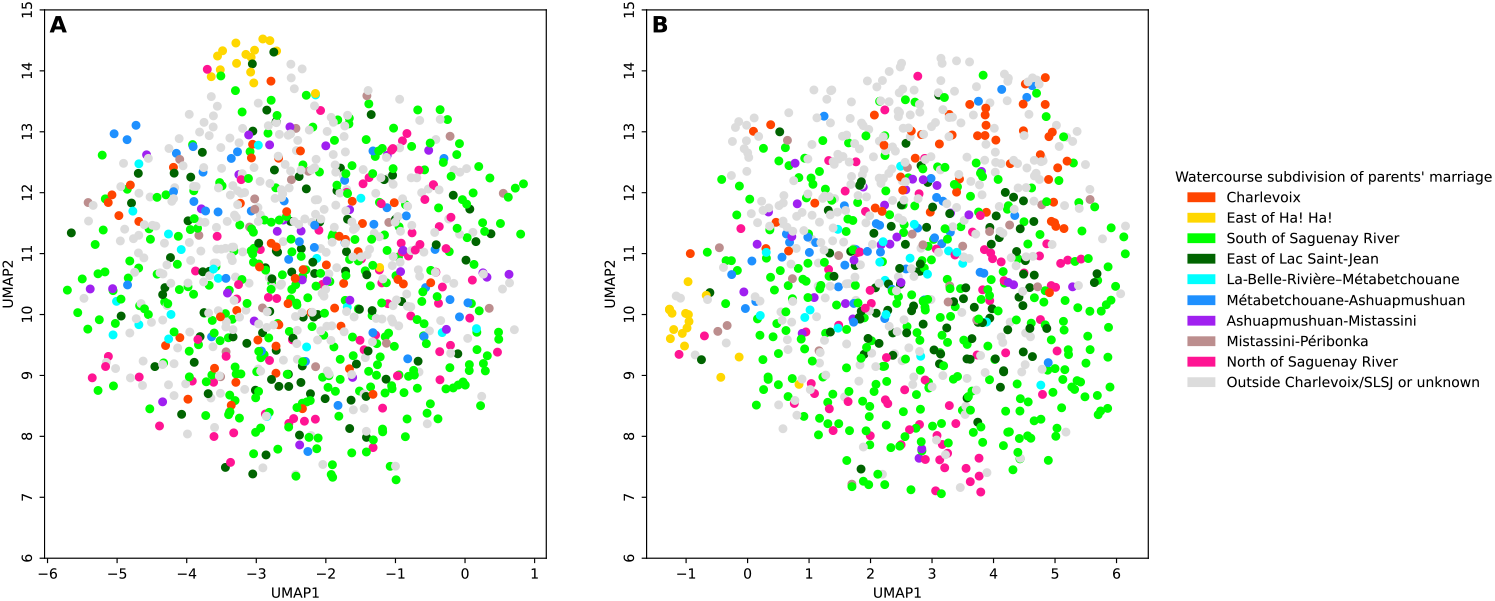
UMAP of CARTaGENE individuals from SLSJ based on pairwise realised (A) and expected (B) kinship. UMAP projection of 904 individuals from the CARTaGENE cohort who are most likely to originate from SLSJ, computed from their (*A*) realised kinship and (*B*) expected kinship transformed as a precomputed distance (*1 – ϕ*). The colours represent the SLSJ watercourse subdivision of parents’ marriage.

### Fine-scale population structure of a whole generation

The great correlation between the population structure observed through genetic and genealogical data^8,21,23,24^ allows us to use the genealogy as a proxy for examining the fine-scale structure of the whole SLSJ population of the last generation available in BALSAC. The UMAP projection of non-siblings probands married between 1931 and 1960 in the SLSJ region reveals concentrations of individuals whose parents were married in the same municipality (Fig. 3). The observed structure follows an east– west gradient, extending from the southeast (East of Ha! Ha! subdivision) to the northwest (Ashuapmushuan–Mistassini subdivision) (see Supplementary Fig. 1). This gradient exhibits a circular pattern centred around a cluster of individuals originating from Charlevoix and concludes with a distinct grouping of individuals whose ancestry lies outside both Charlevoix and SLSJ. More remote, rural municipalities such as L’Anse-Saint-Jean (Supplementary Fig. 5A) exhibit higher levels of concentration in genealogical relationships, while urban centres such as Alma (Supplementary Fig. 5B) display greater dispersion.

**Fig. 3:**
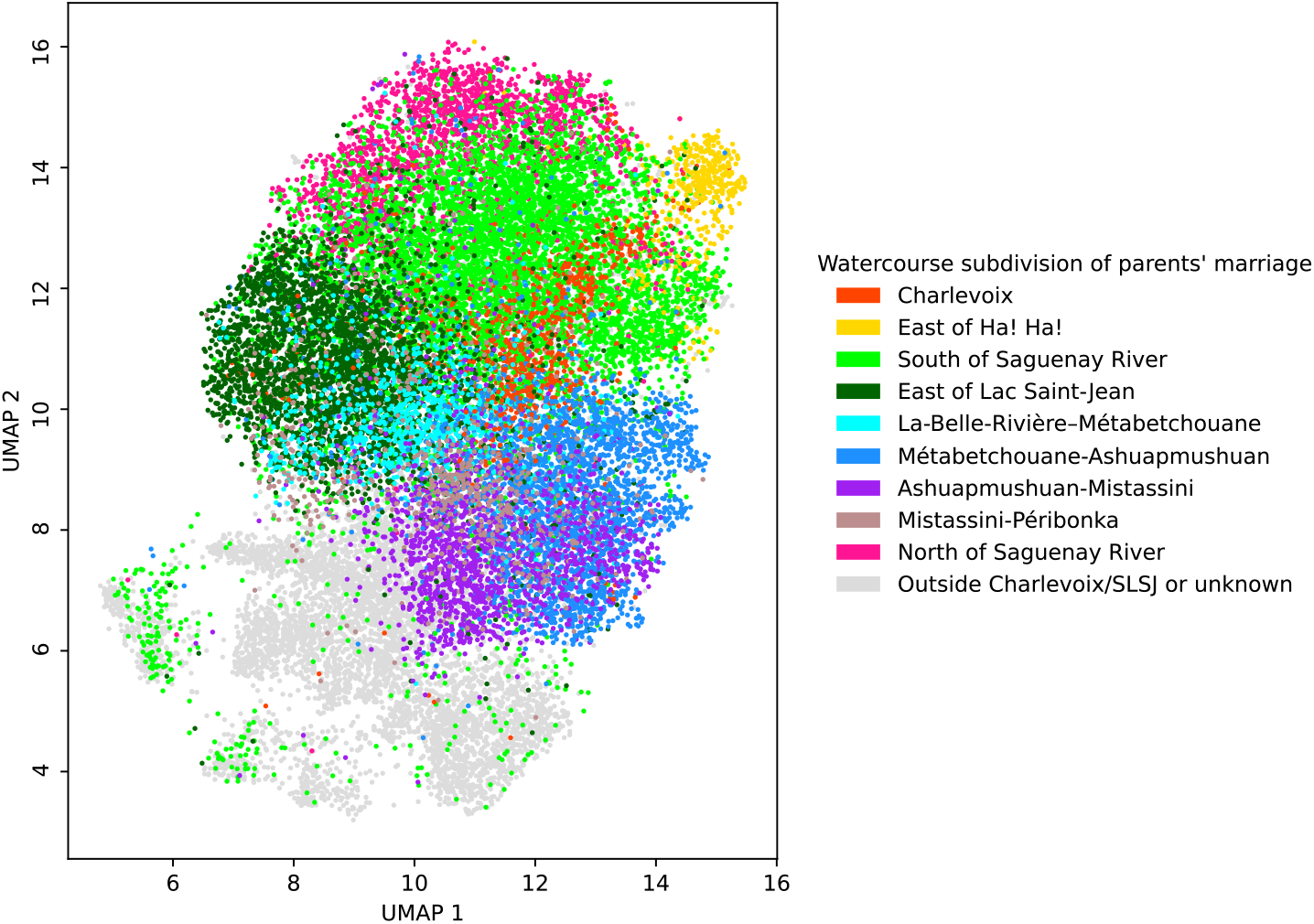
UMAP of pairwise expected kinship for the last-generation SLSJ population. UMAP projection of 26,445 non-siblings who married between 1931 and 1960 in SLSJ, computed from their expected kinship (*ϕ*) transformed as a precomputed distance (1 – *ϕ*). The colours represent the watercourse subdivision of parents’ marriage.

### Expected genetic contribution of founders

To refine our understanding of the migratory patterns underlying the observed structure, we calculated the expected genetic contribution of the SLSJ founders (ancestors married in SLSJ whose parents married elsewhere, see methods) to the SLSJ probands of the last generation. Mean genetic contribution from the SLSJ founders originating from Charlevoix is the highest in the eastern part of SLSJ and progressively declines along an east–west axis (Fig. 4A)^20^. Individuals from the northwesternmost municipalities of Lac-Saint-Jean exhibit more diverse ancestral origins, as well as individuals originating from urban areas, especially south of Saguenay River that exhibit high diversity. A detailed analysis (Fig. 4B) reveals that this east–west genetic gradient is primarily driven by founders from La Malbaie and to some extent Les Éboulements, which both contributed most significantly to the Saguenay area (see also Supplementary Fig. 6). In contrast, Baie-Saint-Paul contributed more to the Lac-Saint-Jean region (Fig. 4B, see also Supplementary Fig. 6). Notably, Les Éboulements contributed disproportionately to the east of Ha! Ha!, a pattern attributable to a wave of migration from approximately 1865 to 1900 that followed an earlier one originating mainly from La Malbaie before 1865 (Supplementary Fig. 6). Notably, east of Ha! Ha! division also presents the highest contribution from Charlevoix founders overall. Individuals from the area between Ashuapmushuan and Mistassini largely trace their ancestry to regions outside Charlevoix (Fig. 4B), with this diversification becoming prominent around 1885 (Supplementary Fig. 7). Namely, contributions from Mauricie are greater in the area between Ashuapmushuan and Mistassini than in other subdivisions (Supplementary Fig. 7). Individuals originating from Gaspésie and Îles-de-la-Madeleine are concentrated in the subdivision south of the Saguenay River (Supplementary Fig. 7).

**Fig. 4:**
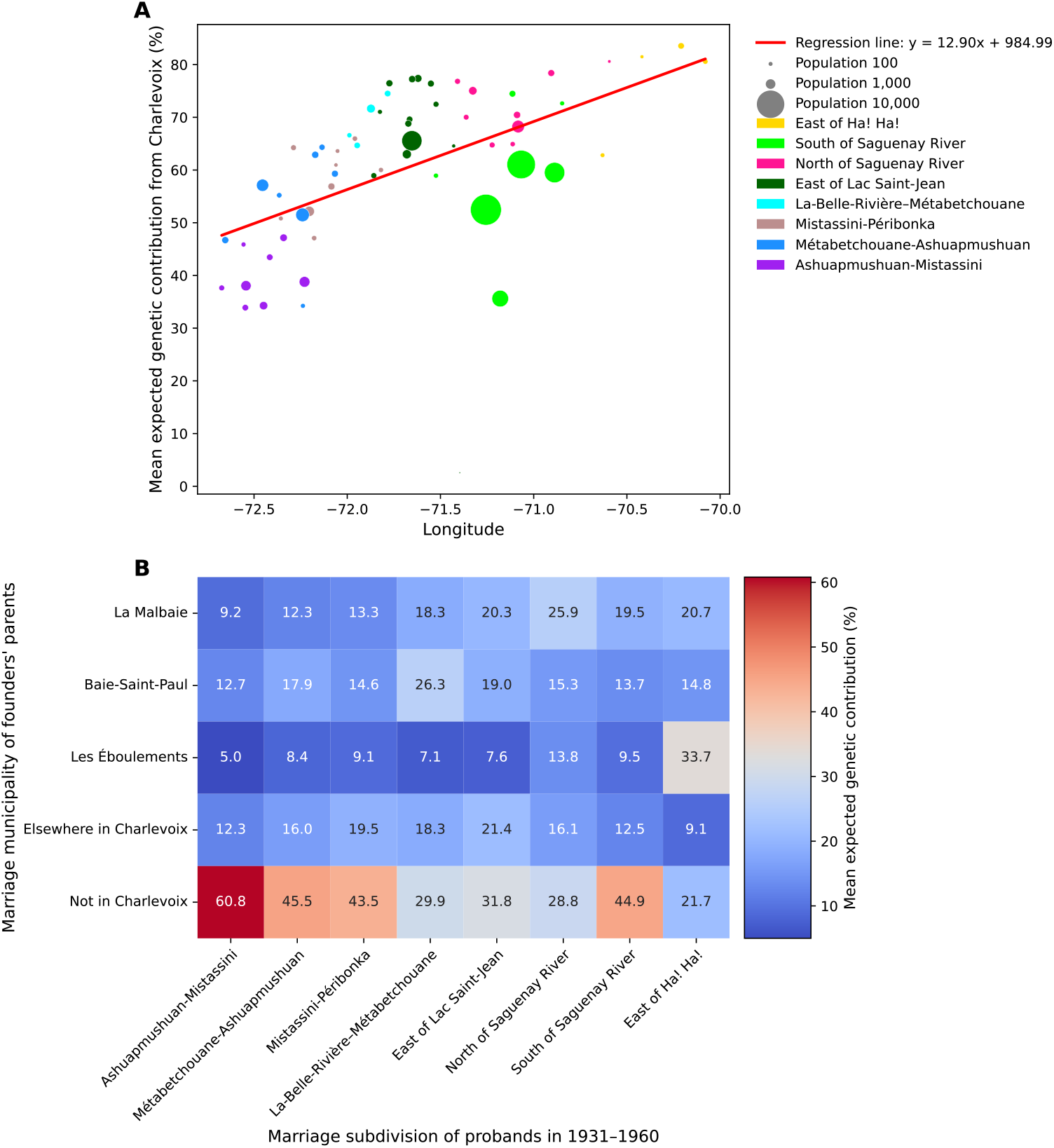
Percentage of the mean expected genetic contribution to probands explained by SLSJ founders depending on geography (A) and on various Charlevoix municipalities (B). Mean expected genetic contribution of SLSJ founders from Charlevoix to probands as a function of the longitude of the probands’ subdivisions (*A*). The red line shows the unweighted linear regression with a Pearson correlation coefficient of 0.517 (*p*-value < 0.001). Mean expected genetic contribution of SLSJ founders from Charlevoix per municipality to probands’ in each subdivision (*B*).

## DISCUSSION

In this study, we found fine-scale population structure in SLSJ, challenging the notion of homogeneity in this founder population^14,18^. The structure is present in both genetic and genealogical data, as previously demonstrated in the more general French-Canadian population^8,21–23^. This fine-scale structure was assessed and visualised on the entire SLSJ population, thanks to deep genealogical data and a time-efficient hybrid algorithm for computing kinship coefficients on such a massive scale.

This fine-scale structure is partly explained by the migratory patterns that shaped the SLSJ population. Previous work described a Charlevoix “gradient,” in which eastern municipalities show a higher genetic contribution from Charlevoix founders than those farther west^20^. However, we show here for the first time at such a finer scale that this gradient is less homogeneous than suggested. Indeed, it centres on La Malbaie, the main contributor to Saguenay, while Baie-Saint-Paul’s influence predominates in Lac-Saint-Jean. In the eastern part of the region, Les Éboulements also shows a strong genetic contribution to the Ha! Ha! Bay area, following an early but previously undocumented wave of migration from La Malbaie. Because Charlevoix families, likely including those from Les Éboulements, often migrated in extended clusters^11^, this family-based movement may explain the steep genetic gradient observed around Ha! Ha!. Moving westward, individuals sampled between Ashuapmushuan and Mistassini show less than half of their genetic input from Charlevoix, instead tracing largely to external Quebec regions other than Charlevoix. In the far northwest of Lac-Saint-Jean, the opening of the Mauricie–Chambord railroad in 1888^9^ brought in diverse founders from outside Charlevoix^20^. By revealing that genetic contributions from Charlevoix vary depending on the SLSJ subdivisions, our work suggests that the distribution of disease-causing variants, mostly originating from Charlevoix, might also be geographically heterogeneous. It is to note that from around 1912, many Acadians settled south of the Saguenay River (particularly in Kénogami), attracted to jobs in the paper and aluminium industries^36^, diluting Charlevoix’s relative genetic contribution in those centres. Likewise, the railroad at Chambord facilitated influx from Quebec regions beyond Charlevoix^20^. These socioeconomic factors further reinforce the heterogeneity of the SLSJ population structure without following strict geographical boundaries, showing that not only geography and migration, but also socioeconomic factors contributed to the gradient and played a role in shaping the structure.

As already known, the fine-scale population structure has an impact on rare variants^3,4^ and association studies^1,2^. Quebec is known for regional enrichments of rare variants, most notably in SLSJ^5^, but evidence points to similar patterns in other less studied regions^4,7,37,38^. Our study suggests that such enrichments may be more localised than at the regional scale, even within regions with a strong founder effect. Within the population of a single region, the carrier rate may vary from one location to another. If a variant is more frequent in one part of the region, and less in another, considering only the overall carrier rate may mitigate the assessment of enrichment and lead to an under-appreciation of the variant’s frequency in the region. This factor is intensified if the carrier tests are performed unevenly, such as in urban centres more than in rural municipalities.

This work showcases some methodological advances, such as the power of parallelised kinship computation via GeneaKit: billions of coefficients inferred efficiently across tens of thousands of individuals. Our hybrid pipeline, melding Karigl’s^25^ and Kirkpatrick’s^27^ algorithms, proved to be time-efficient and is readily extensible to other deep genealogy datasets worldwide. Indeed, whereas computing the kinship coefficients of all SLSJ probands with other software would have been too slow or memory-intensive to be feasible, this task can be achieved in about 2 minutes using 40 CPU cores and 150 GB of RAM. It was not possible to compare directly with Kirkpatrick’s implementation due to the fact that this software is no longer available online, nor with the GENLIB implementation because its execution would require more resources than can be allocated. Following pre-publication of this work, it was brought to our attention that an indirect method by Colleau^39^ can also be applied to compute kinship coefficients at a large scale. Recent work used this approach to compute and visualise all kinship coefficients of a large pedigree of Labrador Retriever^40^, showing the parallel development of kinship computation in human genetics and breeding genetics. By inferring fine-scale population structure using genealogy, we reaffirm the correlation between genetics and genealogy^8,21–23^. Therefore, using genealogy as a proxy for genetic structure is valuable, and it enabled us to include the entire SLSJ population, which would not be possible with genetic data alone, which are unavailable for all individuals, are limited to contemporary generations, and are often concentrated in urban areas. This study also highlights the need for inclusion of smaller and remote communities in future genetic studies, especially for those on rare diseases^41–44^.

While our study provides valuable insights into Quebec’s genetic structure, it is important to acknowledge certain limitations. The CARTaGENE dataset, although rich in genotype data, primarily comprises individuals recruited in urban areas, which might lead to an underrepresentation of more isolated communities. Similarly, the BALSAC genealogies, while extensive, exhibit regional variations in completeness^24^. This uneven data quality could potentially introduce biases when detecting population structure. Furthermore, our use of parents’ marriage location appears to be a better proxy for regional affiliation than the recruitment locations of CARTaGENE. It may still not fully account for individual or family mobility over generations or situations where parents originate from different regions. Despite these limitations, our rigorous quality control, which involved filtering out individuals with more than half null kinship coefficients, allowed us to observe the Quebec-wide genetic structure previously described. This resolution was sufficient to distinguish the peripheries of the SLSJ region. We also found that our UMAP projections, despite the inherent stochastic nature of the algorithm that can make direct comparisons challenging^34^, yielded highly similar results with matrices of expected and realized kinship. These findings underscore the robustness of our analysis in capturing the underlying genetic landscape of SLSJ and the province of Quebec.

In conclusion, integrating large-scale genealogical and genotype data reveals fine-scale structure in SLSJ that broadly aligns with geographic gradients: eastern municipalities bear stronger Charlevoix ancestry, western ones show increased contribution from other Quebec regions, shaped in part by historic transport routes. Crucially, this gradient is not uniform within Charlevoix, our analysis reveals a distinct and previously under-appreciated migratory stream from Les Éboulements to the east of Ha! Ha!. Such heterogeneity suggests uneven distributions of founder variants and, by extension, locally variable prevalence of associated genetic diseases. These findings also have implications for how we understand human genetic diversity more broadly. The use of self-reported race and ethnicity as proxies for human genetic diversity has long been widespread, and recent applications of UMAP have been employed to support such categorisations^45^. In reality, population structure and genetic diversity are observable not only at the continental scale, but also within regional populations^46–48^, such as those within the province of Quebec. Our findings further demonstrate that population structure can be detected at even finer geographic resolutions, down to the level of municipalities, where we observe gradients of admixture in addition to sharply defined clusters for more remote localities. Such subcontinental gradients can impact genetic association studies, and should not be overlooked^49,50^.

## DATA AVAILABILITY

Quebec genotype data is available via an independent data access committee by the CARTaGENE cohort (https://cartagene.qc.ca/en/researchers/access-request.html). Quebec genealogical data is available upon request to BALSAC (https://balsac.uqac.ca). These datasets are under restricted access due to the informed consent given by study participants.

## CODE AVAILABILITY

The GeneaKit library is available under the open-source MIT licence in the following GitHub repository: https://github.com/Genopop/geneakit. The code used for this study can be found in the following GitHub repository: https://github.com/Genopop/slsj_structure.

## ACKNOWLEDGEMENTS

This work was made possible by the Digital Research Alliance of Canada which provided access to storage and computing resources. We are extremely grateful to all participants in this research.

## AUTHOR CONTRIBUTIONS

All authors acquired data and approved the final version of the manuscript. GPM and CM played an important role in interpreting the results. GPM, CM and SLG conceived and designed the study and drafted the manuscript. ADB revised the manuscript.

## ETHICAL APPROVAL

This study was approved by the ethics board of the Université du Québec à Chicoutimi (UQAC).

