## Supplementary Material for "Fine-scale structure of a whole regional population through genetics and genealogies"

**Supplementary Table 1: Municipalities of watercourse-defined subdivisions of Saguenay–Lac-Saint-Jean (SLSJ).**

| **Subdivision (No of municipalities)** | Municipalities |
|---|---|
| **East of Ha! Ha!** (4) | Petit-Saguenay, L'Anse-Saint-Jean, Rivière-Éternité, Saint-Félix-d'Otis |
| **South of Saguenay River** (9) | Ferland-et-Boilleau, La Baie, Chicoutimi, Laterrière, Arvida, Jonquière, Saint-Ambroise, Bégin, Larouche |
| **East of Lac Saint-Jean** (13) | Mont-Apica, Notre-Dame-du-Rosaire, Labrecque, Saint-Nazaire, Delisle, Saint-Bruno, Alma, Hébertville-Station, L'Ascension-de-Notre-Seigneur, Hébertville, Saint-Gédéon, Saint-Henri-de-Taillon, Sainte-Monique-de-Honfleur |
| **La-Belle-Rivière–Métabetchouane** (4) | Lac-à-la-Croix, Métabetchouan, Desbiens, Saint-André-du-Lac-Saint-Jean |
| **Métabetchouane-Ashuapmushuan** (7) | Saint-François-de-Sales, Lac-Bouchette, Mashteuiatsh, Roberval, Sainte-Hedwidge, Saint-Félicien, La Doré |
| **Ashuapmushuan-Mistassini** (8) | Dolbeau, Saint-Prime, Saint-Méthode, Albanel, Normandin, Girardville, Saint-Edmond, Saint-Thomas-Didyme |
| **Mistassini-Péribonka** (11) | Chute-des-Passes, Saint-Ludger-de-Milot, Saint-Augustin-du-Lac-Saint-Jean, Péribonka, Sainte-Élisabeth-de-Proulx, Chambord, Sainte-Jeanne-d'Arc-du-Lac-Saint-Jean, Saint-Stanislas-du-Lac-Saint-Jean, Mistassini, Saint-Eugène-du-Lac-Saint-Jean, Notre-Dame-de-Lorette |
| **North of Saguenay River** (7) | Sainte-Rose-du-Nord, Saint-Fulgence, Chicoutimi-Nord, Saint-Honoré-de-Chicoutimi, Saint-David-de-Falardeau, Shipshaw, Saint-Charles-de-Bourget |

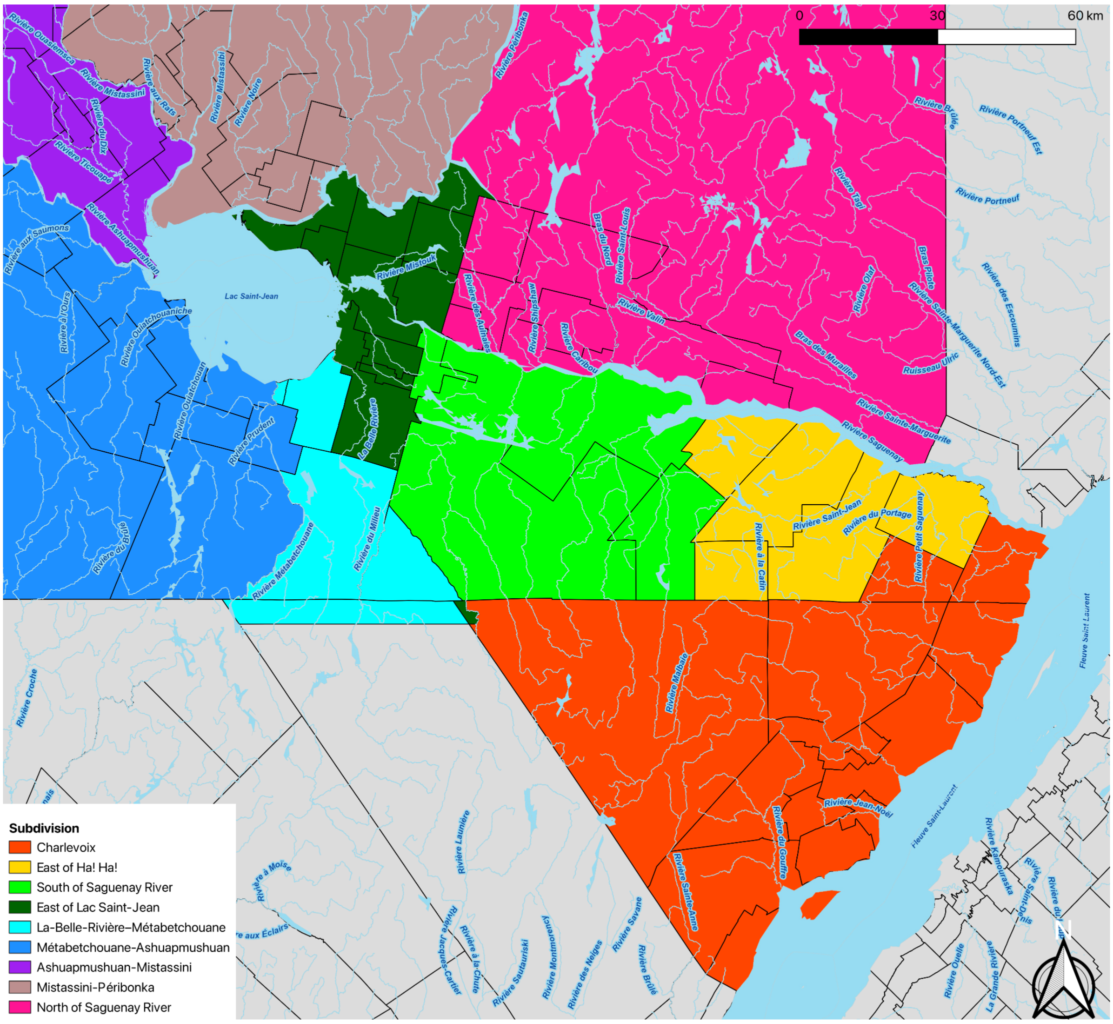


**Supplementary Fig. 1: Watercourse-defined subdivisions of Saguenay–Lac-Saint-Jean and Charlevoix.**

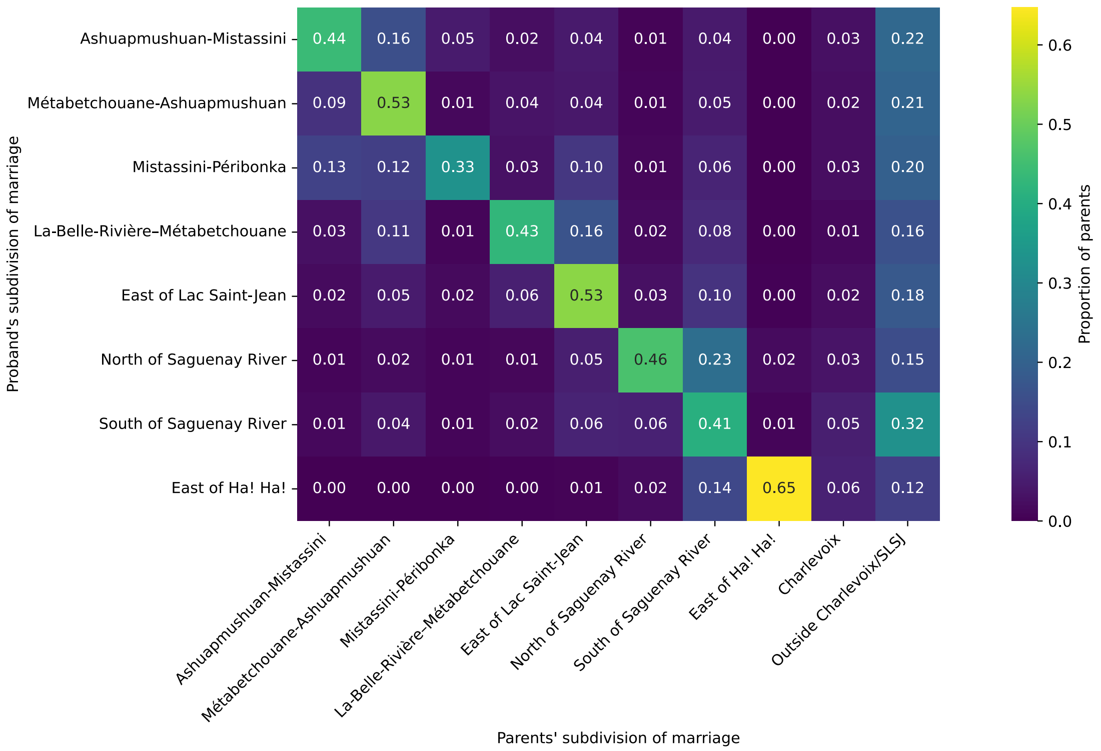


**Supplementary Fig. 2: Migration within and between subdivisions.** Proportion of parents married in each watercourse subdivision (column) for each SLSJ probands' subdivision of marriage (row). The sum of each row is equal to one.

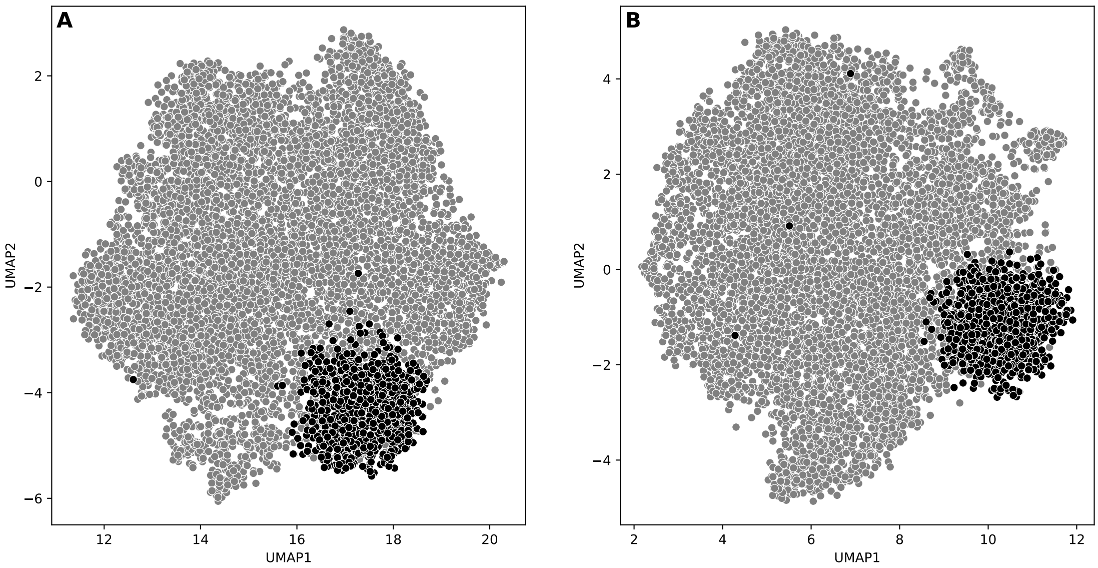


**Supplementary Fig. 3: HDBSCAN clustering of CARTaGENE individuals most likely originating from SLSJ.** (*A*) UMAP projection of 7,970 individuals from the CARTaGENE cohort, computed from their (*A*) realised kinship and (*B*) expected kinship transformed as a precomputed distance ($1 - \phi$). Individuals most likely originating from the SLSJ, as identified by HDBSCAN clustering, are shown in black.

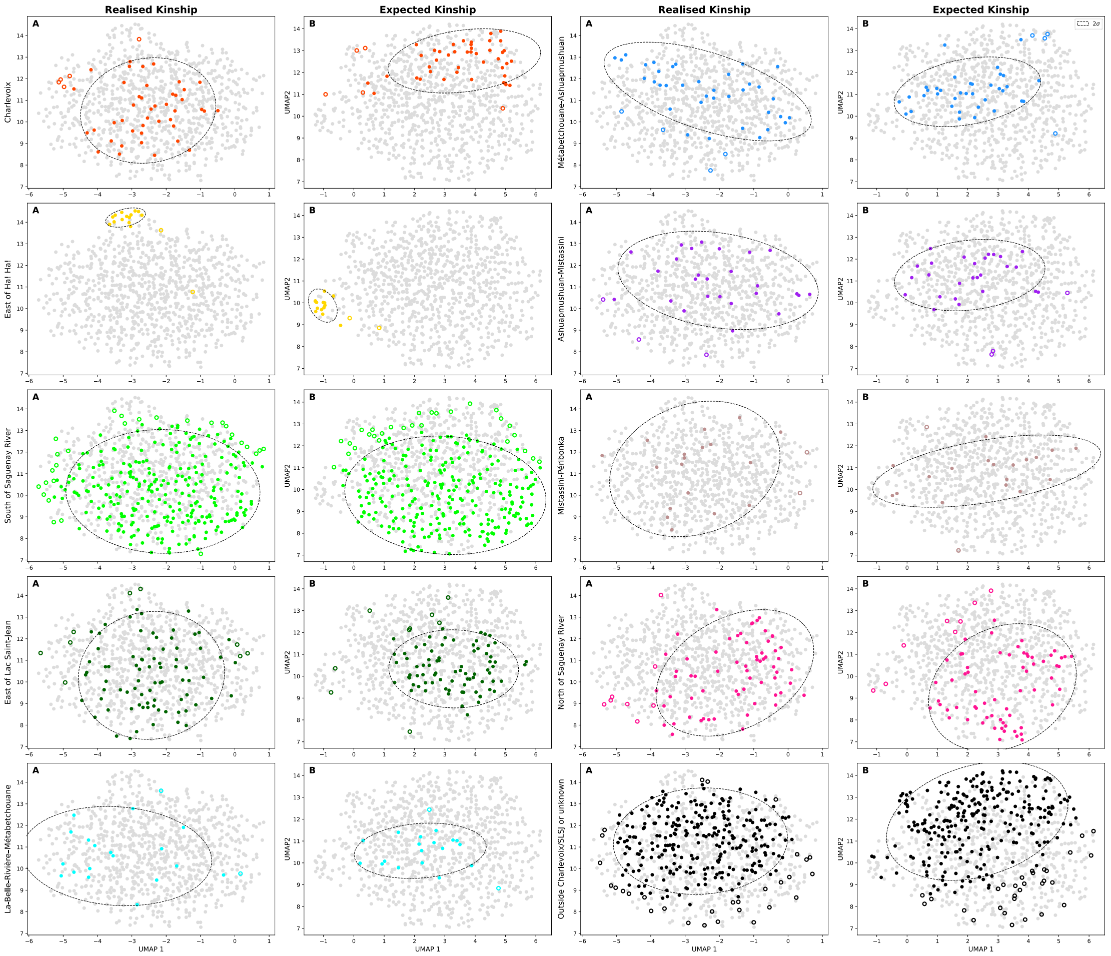


**Supplementary Fig. 4: Distribution of CARTaGENE SLSJ individuals whose parents married in each watercourse subdivision.** UMAP projection of 904 individuals from the CARTaGENE cohort most likely originating from SLSJ, computed from their (*A*) realised kinship and (*B*) expected kinship transformed as a precomputed distance ($1 - \phi$). The coloured dots represent individuals whose parents married in the indicated subdivision of SLSJ. Open circles are individuals who were identified as outliers using an ellipse learned from their Gaussian distribution. The confidence ellipses of the remaining inliers have a radius of two standard deviations.

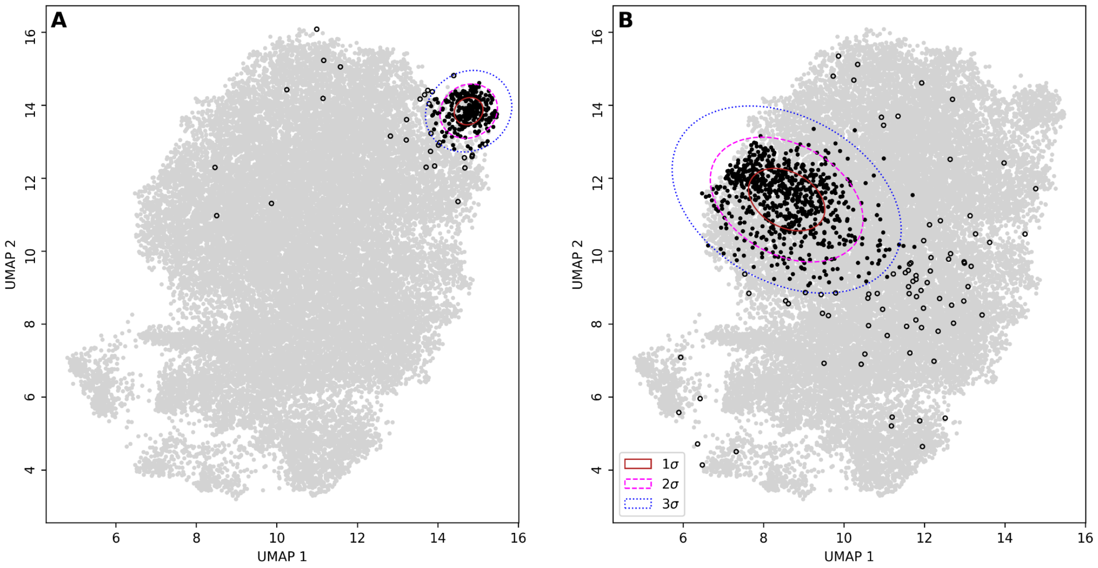


**Supplementary Fig. 5: Difference in the spatial dispersion of individuals between a rural (A) and an urban (B) municipality.** UMAP projection of 26,445 non-siblings who married between 1931 and 1960 in SLSJ, computed from their expected kinship ($\phi$) transformed as a precomputed distance ($1 - \phi$). Black dots represent individuals whose parents married in A) L'Anse-Saint-Jean; and *B)* Alma. Black open circles are individuals who were identified as outliers, whereas the confidence ellipses of the remaining inliers have a radius of one, two, and three standard deviations.

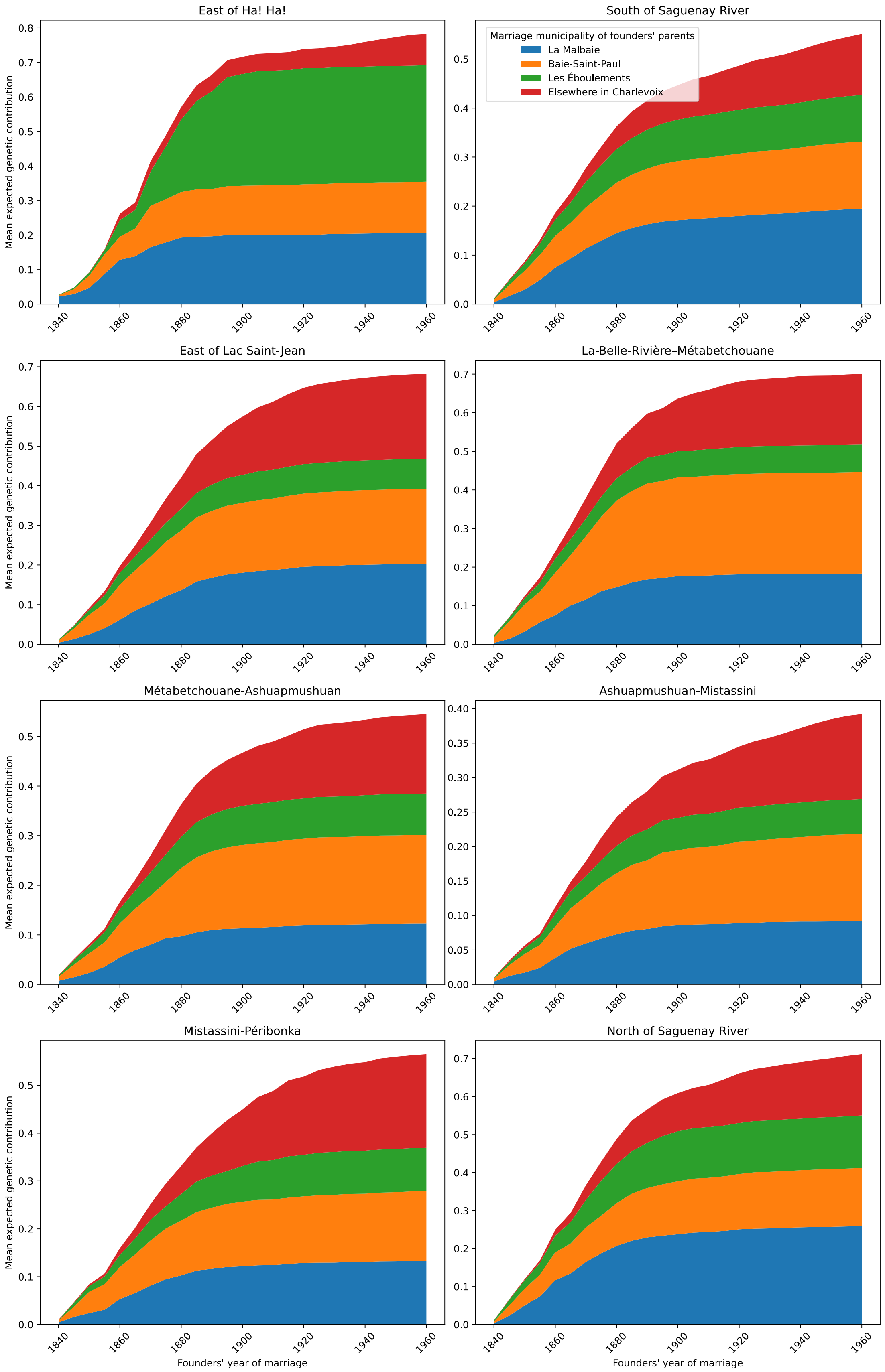


**Supplementary Fig. 6: Cumulative genetic contribution of SLSJ founders originating from Charlevoix to probands in each watercourse subdivision per period of arrival.**

Cumulative mean expected genetic contribution of SLSJ founders per municipality of Charlevoix and period of marriage to probands married in various subdivisions.

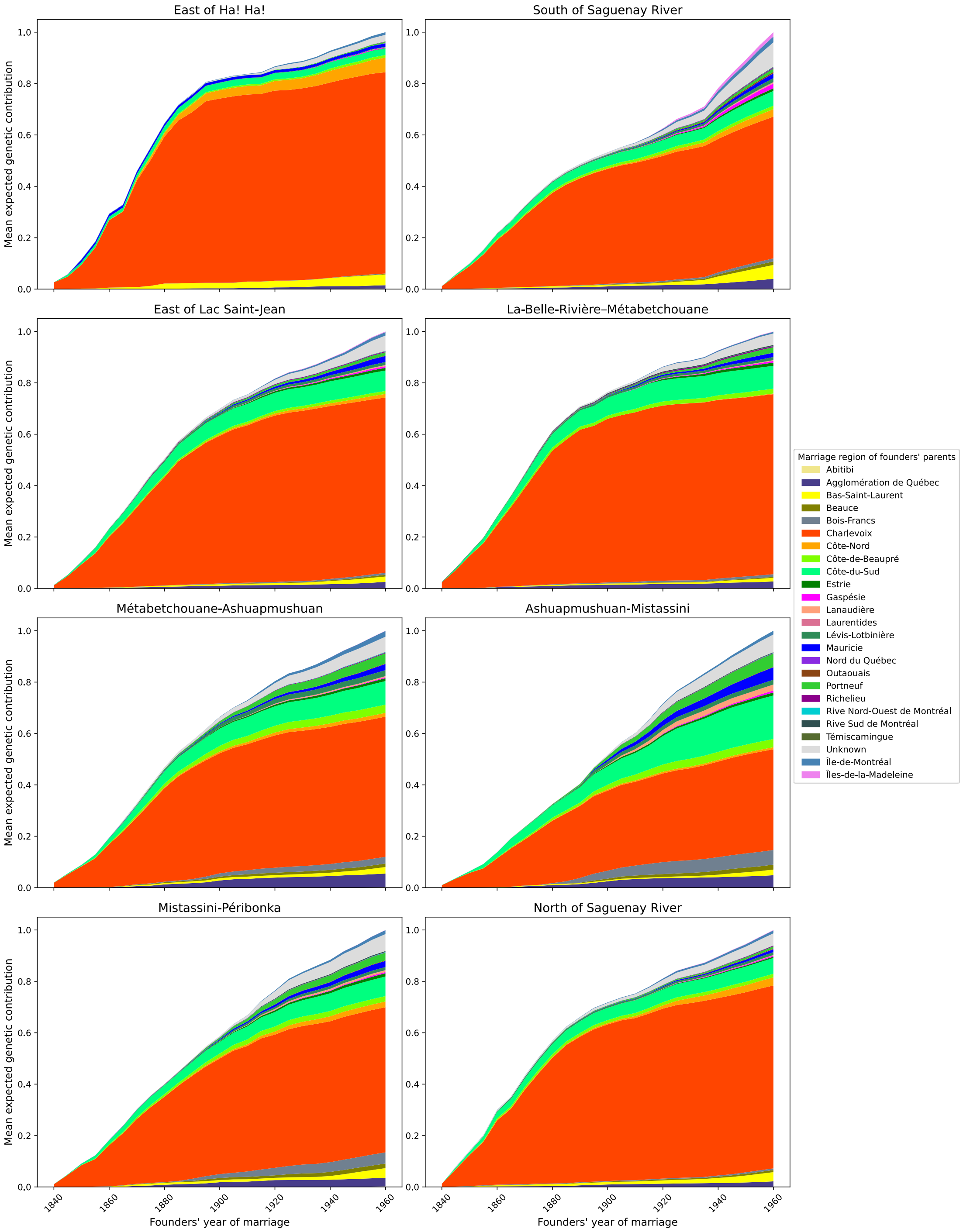


**Supplementary Fig. 7: Cumulative genetic contribution of SLSJ founders originating from Quebec to probands in each watercourse subdivision per period of arrival.** Cumulative mean expected genetic contribution of SLSJ founders per Quebec regions and period of marriage to probands married in various SLSJ subdivisions.

**Supplementary Text 1: Pseudocode for novel kinship algorithm.** In the C++ implementation of this algorithm, *genealogy* is a hash map of a data structure named *individual*, referenced through a unique integer ID. The genealogy is sorted so that any ancestor appears before their offspring. Each *individual* contains notably a reference to two other *individual* structures, a father (if present in the genealogy) and a mother (if present). It also possesses an immutable *rank* from 1 to *n* individuals in the genealogy, which indicates the individual's order in the hash map. Finally, its mutable *founder_index* indicates whether the individual is currently a founder (if non null) and, if that's the case, where their kinship values are located in the current founder kinship matrix. The function `compute_kinships(`*`genealogy, proband_IDs`*`)` is called using the hash map and a list of the probands' integer IDs, and returns a kinship matrix with the probands in the same order as in the list. The function `compute_kinship_between_probands(probands, founder_matrix, proband_matrix)` may run the nested for loop in parallel, for each generation.

```
FUNCTION get_previous_generation(genealogy, current_individuals):
    CREATE a new empty set called 'parents'
    FOR each ID in 'current_individuals':
        GET the person from 'genealogy' using the ID
        IF person has a father:
            ADD father's ID to 'parents'
        IF person has a mother:
            ADD mother's ID to 'parents'
    CREATE a new list called 'previous_generation' from the unique IDs in 'parents'
    SORT 'previous_generation' by ID
    RETURN 'previous_generation'

FUNCTION get_generations(genealogy, starting_probands):
    CREATE an empty list of lists called 'generations'
    CREATE an empty list called 'current_generation'
    FOR each ID in starting_probands:
        ADD ID to 'current_generation'
    WHILE 'current_generation' is not empty:
        ADD 'current_generation' to 'generations'
        SET 'current_generation' to the result of calling
get_previous_generation(genealogy, current_generation)
    RETURN 'generations'

FUNCTION copy_bottom_up(generations):
    CREATE an empty list of lists called 'bottom_up'
    CREATE an empty list called 'first_generation'
    FOR each ID in the first list of 'generations':
        ADD ID to 'first_generation'
    ADD 'first_generation' to 'bottom_up'
    FOR each index 'i' from the first to the second-to-last list of 'generations':
        CREATE an empty list called 'combined_generation'
        CREATE a set called 'previous_generation' from the individuals in bottom_up[i]
        CREATE an empty set called 'next_generation'
        FOR each ID in generations[i + 1]:
            ADD ID to 'next_generation'
        PERFORM a set union of 'previous_generation' and 'next_generation', adding the
unique results to 'combined_generation'
        ADD 'combined_generation' to 'bottom_up'
```

```
    REVERSE 'bottom_up' to go from oldest to youngest
    RETURN 'bottom_up'

FUNCTION copy_top_down(generations):
    REVERSE the order of 'generations'
    CREATE an empty list of lists called 'top_down'
    CREATE an empty list called 'first_generation'
    FOR each ID in the first list of the (now reversed) 'generations':
        ADD ID to 'first_generation'
    SORT 'first_generation'
    ADD 'first_generation' to 'top_down'
    FOR each index 'i' from the first to the second-to-last list of 'generations':
        CREATE an empty list called 'combined_generation'
        CREATE an empty set called 'previous_generation'
        FOR each ID in top_down[i]:
            ADD ID to 'previous_generation'
        CREATE an empty set called 'next_generation'
        FOR each ID in generations[i + 1]:
            ADD ID to 'next_generation'
        PERFORM a set union of 'previous_generation' and 'next_generation', adding the
unique results to 'combined_generation'
        ADD 'combined_generation' to 'top_down'
    RETURN 'top_down'

FUNCTION intersect_both_directions(bottom_up, top_down):
    CREATE an empty list of lists called 'vertex_cuts'
    FOR each index 'i' for every list in 'bottom_up':
        CREATE an empty list called 'current_vertex_cut'
        CREATE a set called 'set_from_bottom_up' from the IDs in bottom_up[i]
        CREATE a set called 'set_from_top_down' from the IDs in top_down[i]
        CREATE an empty list called 'common_individuals'
        PERFORM a set intersection of 'set_from_bottom_up' and 'set_from_top_down',
adding the unique results to 'common_individuals'
        FOR each ID in 'common_individuals':
            ADD ID to 'current_vertex_cut'
        ADD 'current_vertex_cut' to 'vertex_cuts'
    RETURN 'vertex_cuts'

FUNCTION cut_vertices(genealogy, proband_IDs):
    CREATE empty lists of lists called 'generations' and 'vertex_cuts'
    SET 'generations' to the result of calling get_generations(genealogy, proband_IDs)
    CREATE empty lists of lists called 'bottom_up' and 'top_down'
    SET 'bottom_up' to the result of calling copy_bottom_up(generations)
    SET 'top_down' to the result of calling copy_top_down(generations)
    SET 'vertex_cuts' to the result of calling intersect_both_directions(bottom_up,
top_down)
    SET the last list in 'vertex_cuts' to 'proband_IDs'
    RETURN 'vertex_cuts'

FUNCTION compute_kinship(ind1, ind2, founder_matrix):
    SET 'kinship' to 0.0
    SET 'founder_index1' to individual ind1's founder_index
    SET 'founder_index2' to individual ind2's founder_index
    IF 'founder_index1' is not null AND 'founder_index2' is not null:
        SET 'kinship' to founder_matrix[founder_index1][founder_index2]
    ELSE IF 'founder_index1' is not null:
        IF ind2 has a father:
            ADD 0.5 * compute_kinship(ind1, ind2's father, founder_matrix) to
```

```
'kinship'
        IF ind2 has a mother:
            ADD 0.5 * compute_kinship(ind1, ind2's mother, founder_matrix) to
'kinship'
    ELSE IF 'founder_index2' is not null:
        IF ind1 has a father:
            ADD 0.5 * compute_kinship(ind1's father, ind2, founder_matrix) to
'kinship'
        IF ind1 has a mother:
            ADD 0.5 * compute_kinship(ind1's mother, ind2, founder_matrix) to
'kinship'
    ELSE IF ind1's rank is equal to ind2's rank:
        SET 'kinship' to 0.5
        IF ind1 has a father AND ind2 has a mother:
            ADD 0.5 * compute_kinship(ind1's father, ind2's mother, founder_matrix) to
'kinship'
    ELSE IF ind1's rank is less than ind2's rank:
        IF ind2 has a father:
            ADD 0.5 * compute_kinship(ind1, ind2's father, founder_matrix) to
'kinship'
        IF ind2 has a mother:
            ADD 0.5 * compute_kinship(ind1, ind2's mother, founder_matrix) to
'kinship'
    ELSE:
        IF ind1 has a father:
            ADD 0.5 * compute_kinship(ind1's father, ind2, founder_matrix) to
'kinship'
        IF ind1 has a mother:
            ADD 0.5 * compute_kinship(ind1's mother, ind2, founder_matrix) to
'kinship'
    RETURN 'kinship'

FUNCTION compute_kinship_with_oneself(probands, founder_matrix, proband_matrix):
    FOR each index 'i' for every proband in 'probands':
        SET 'proband' to probands[i]
        SET 'kinship' to 0.5
        IF proband has a father AND proband has a mother:
            ADD 0.5 * compute_kinship(proband's father, proband's mother,
founder_matrix) to 'kinship'
        SET proband_matrix[i][i] to 'kinship'

FUNCTION compute_kinship_between_probands(probands, founder_matrix, proband_matrix):
    FOR each index 'i' for every proband in 'probands':
        SET 'proband1' to probands[i]
        FOR each index 'j' from the first proband up to (but not including) the
current index 'i':
            SET 'proband2' to probands[j]
            SET 'kinship' to compute_kinship(proband1, proband2, founder_matrix)
            SET proband_matrix[i][j] to 'kinship'
            SET proband_matrix[j][i] to 'kinship'

FUNCTION compute_kinships(genealogy, proband_IDs):
    SET 'vertex_cuts' to the result of calling cut_vertices(genealogy, proband_IDs)
    CREATE a new matrix called 'current_founder_matrix' with dimensions (size of
vertex_cuts[1], size of vertex_cuts[1]) and filled with zeros
    FOR each index 'i' for every individual in the first vertex cut:
        SET current_founder_matrix[i][i] to 0.5
    FOR each index 'i' from the first to the second-to-last vertex cut:
```

```
        SET 'founder_index' to 1
        FOR each person ID in vertex_cuts[i]:
            GET the person from the genealogy using their ID
                SET person's founder index (null by default) to 'founder_index'
            INCREMENT 'founder_index'
        CREATE a new matrix called 'current_proband_matrix' with dimensions (size of
vertex_cuts[i + 1], size of vertex_cuts[i + 1])
        CREATE an empty list of persons called 'current_probands'
        FOR each person ID in vertex_cuts[i + 1]:
            ADD 'person' from 'genealogy' using their ID to 'current_probands'
        CALL compute_kinship_with_oneself(current_probands, current_founder_matrix,
current_proband_matrix)
        CALL compute_kinship_between_probands(current_probands,
current_founder_matrix, current_proband_matrix)
        SET 'current_founder_matrix' to 'current_proband_matrix'
    RETURN 'current_founder_matrix'
```
